# Iron exposure reprograms redox homeostasis through mitochondrial depolarization in neuroblastoma

**DOI:** 10.64898/2026.10.08.755735

**Authors:** Ana Paula M. Mendonça, Stephane Claverol, Bernardo F. Braz, Aline S. Freire, Franklin D. Rumjanek, Ricardo E. Santelli, Caroline Tokarski, Danielle B. Carvalho, Rodrigue Rossignol, Nivea D. Amoedo, Marcus F. Oliveira

## Abstract

**Purpose:** Neuroblastoma is the most common malignancy of infancy, and cancer cells strongly depend on iron metabolism for proliferation. Whether sub-cytotoxic iron load reprograms mitochondrial and redox metabolism, and how this relates to the malignant behavior of neuroblastoma cells, is poorly defined. We hypothesized that exposure of human neuroblastoma cells to a non-toxic iron overload perturbs mitochondrial and redox metabolism, thereby affecting cancer cell properties.

**Methods:** SH-SY5Y neuroblastoma cells were exposed to 100 µM iron sulfate for 24 h. The resulting changes were monitored using complementary methods, including cell viability, cytotoxicity, proliferation and migration assays, intracellular iron quantification by ICP-MS, confocal imaging of mitochondrial morphology and membrane potential (Δψ_m_), high-resolution respirometry with flux control ratios, probe-based oxidant and H_2_O_2_ measurements, and label-free quantitative proteomics with Ingenuity Pathway Analysis.

**Results:** Non-toxic iron loading raised intracellular iron about four-fold, promoted cell proliferation and reduced migration. Iron induced mitochondrial fragmentation and Δψ_m_ depolarization while preserving mitochondrial content and OXPHOS coupling efficiency. Among respiratory states, only proton-leak respiration increased, paralleled by a higher leak/ETS coupling ratio. Total cellular oxidant levels and H_2_O_2_ output were strongly reduced. The proteome showed downregulation of ribosomal proteins and translation/RNA-processing components and upregulation of histones and ferritin.

**Conclusion:** Mild iron exposure reprograms mitochondrial and redox metabolism in neuroblastoma cells. The observed selective increase in proton leak lowers Δψ_m_ to a range that biophysically limits oxidant production while OXPHOS coupling is preserved. Interfering with this adaptive, proton leak-dependent and preventive antioxidant program may offer a way to alter neuroblastoma progression.

## 1. Introduction

Pediatric cancers are heterogeneous malignancies of children and adolescents under 20 years of age, and, despite their relatively low incidence, cancer is the second leading cause of death among children aged 1–14 years in the United States [1]. Neuroblastoma, the most common cancer diagnosed in the first year of life, arises from sympathetic nervous system progenitors derived from embryonic neural crest cells and shows a wide clinical spectrum, from low-risk tumors that may regress spontaneously to high-risk metastatic disease with poor outcomes [2]. Primary tumors usually originate in the adrenal glands, but high-risk patients often present with metastases in lymph nodes, liver, bone and bone marrow, and metastatic disease correlates strongly with poor prognosis [3]. *MYCN* amplification is a hallmark of neuroblastoma aggressiveness [4], and recurrent alterations are also found in *ALK* [5], *ATRX* [6] and *TERT* [7–9]. Despite more specific therapies, mortality remains high in high-risk patients [10, 11], and the biochemical and metabolic bases of neuroblastoma progression remain largely unknown.

Altered energy metabolism is a recognized hallmark of cancer [12]. Tumor energy metabolism is heterogeneous: whereas some tumors display enhanced aerobic glycolysis and become highly glucose-dependent, so-called ‘high-OXPHOS’ tumors rely primarily on mitochondrial oxidative phosphorylation to sustain growth and survival [13]. Yet, the changes in neuroblastoma cells remain poorly described. Intra-operative [U-^13^C]glucose infusion has shown that neuroblastomas exhibit the highest lactate labelling among childhood cancers and elevated tyrosine-catabolism metabolites, indicating increased catecholamine production [14]. The transcription factor *MYCN* promotes the high-OXPHOS subtype of neuroblastoma [15]; *MYCN*-overexpressing cells are highly proliferative and rely on mitochondrial lipid oxidation, rendering them sensitive to β-oxidation inhibition [16, 17].

Iron is a redox-cycling transition metal whose chemistry makes it an excellent cofactor for enzymatic electron transfer [18]. Protein-bound iron is essential for oxygen transport, oxidant detoxification, and energy metabolism [19], but in its labile, “free” state it is a powerful pro-oxidant that catalyzes metal-mediated oxidation of enzymes [20] and generates a range of cytotoxic oxidants and free radicals [21]. Epidemiological data links relatively high body iron to increased cancer risk [22, 23]. Although the mechanisms are incomplete, tumors commonly restrict iron bioavailability at the systemic level [24, 25] while, at the cellular level, cancer cells favor iron uptake and retention to expand the labile iron pool and foster proliferation [26–28]. Dysregulated iron homeostasis is also seen in neuroblastoma, where functional iron depletion and elevated serum ferritin are associated with poor outcome [25, 29–31], and neuroblastoma cells induce ferritin, remain viable and proliferate even under severe iron overload [32–37]. Consistently, pharmacological iron chelation has antitumor effects in neuroblastoma [38, 39]. Mild iron overload elicits adaptive mitochondrial responses, including a modest proton leak without apparent loss of bioenergetic efficiency or ATP levels [40], whereas iron-store depletion through chelation promotes mitophagy and reduces mitochondrial content [41]. Together, these observations suggest mitochondrial metabolic flexibility under mild iron loading, plausibly to limit redox damage and preserve viability.

Because neuroblastoma cells strongly depend on iron for viability and proliferation [32–35, 37, 42], we hypothesized that a mild, sub-cytotoxic iron overload would selectively increase mitochondrial proton leak and lower cellular oxidant output, remodeling cell behavior without compromising OXPHOS coupling. We deliberately frame the study as cell-autonomous metabolic reprogramming driven by a micronutrient cue (iron), recognizing that iron availability is itself a microenvironmental and systemic variable in tumors. Evaluating this hypothesis at cellular, biochemical and proteomic levels in a human neuroblastoma cell line, we find that mild iron exposure promotes proliferation while reducing migration, increases proton-leak respiration with preserved coupling, and strongly suppresses cellular oxidants, in association with mitochondrial structural and functional remodeling.

## 2. Materials and methods

### 2.1. Cell culture and authentication

Human neuroblastoma SH-SY5Y cells were obtained from the Rio de Janeiro Cell Bank (BCRJ) and cultured at 37 °C in a humidified atmosphere of 95 % air and 5 % CO_2_. Cells were maintained in Minimum Essential Medium (MEM; Thermo Scientific, USA) supplemented with 10 % fetal bovine serum (FBS; LGCBio, Brazil), 10 U/mL penicillin–streptomycin (Thermo Scientific), 0.5 µg/mL polymyxin B sulfate (Sigma-Aldrich, USA) and Ham’s F-12 Nutrient Mix (1:1) (Thermo Scientific). Cells were used up to passage 21, subcultured 1:20 in 25 cm^2^ flasks every 3 days. The identity of the SH-SY5Y line (a non-MYCN-amplified neuroblastoma line) and its authentication record were as provided by BCRJ; cells were routinely checked for the absence of mycoplasma contamination.

### 2.2. Iron treatment

Iron sulfate heptahydrate (Sigma-Aldrich) was freshly prepared before each experiment in distilled water at a 1.4 mg/mL stock, sterilized through a 0.22-µm PES filter. At ∼90 % confluence, cultures were washed twice with PBS, the medium was replaced with fresh medium containing iron sulfate at the indicated concentrations, and cells were incubated for 24 h at 37 °C and 5 % CO_2_. A concentration of 100 µM iron sulfate for 24 h was used as the standard non-toxic condition (see Results).

### 2.3. Protein quantification

Protein content was determined by the Lowry method with bovine serum albumin as standard [43].

### 2.4. Cellular iron quantification

Intracellular iron was determined by inductively coupled plasma mass spectrometry (ICP-MS). Cells grown in 90-cm^2^ dishes (0.5 × 10^6^ cells/well) and treated or not with iron were trypsinized, centrifuged at 321 × g for 10 min at 21.5 °C, and resuspended in 2 mL FBS-free MEM-F12. Samples were diluted 1:6 or 1:15 (v/v) in ultrapure water (Milli-Q Direct 8; Merck Millipore, USA). Analyte concentrations were obtained by interpolation against seven-point calibration curves prepared from a 1000 mg/L spectral standard (Quimlab Chemistry & Metrology, Brazil). Measurements used a Thermo Fisher Scientific iCAP Qc ICP-MS (Bremen, Germany) equipped with a collision/reaction cell, nickel sample and skimmer cones, a quartz double-pass cyclonic spray chamber, a Teflon concentric nebulizer (FPA-ST), Peltier cooling, a 2.5-mm quartz injector, and a CETAC ASX-520 autosampler (CETAC Technologies, USA); Qtegra 1.5.1189.1 software was used for acquisition. Operating conditions were: incident power 1550 W; plasma gas 14 L/min; auxiliary gas 0.80 L/min; nebulizer gas 1.0 L/min; dwell time 10 ms; one channel per mass unit. Iron was measured as ^54^Fe, ^56^Fe and ^57^Fe in kinetic energy discrimination (KED) mode using helium at 5.025 mL/min. Transport and ionization interferences were corrected by internal standardization with ^45^Sc.

### 2.5. Cellular viability

Viability was assessed by MTT reduction. Cells (0.1 × 10^6^/well, 96-well plates) were incubated for 24 h with iron sulfate. A death control used 20 % DMSO (v/v) for 24 h. After incubation, the medium was removed, cells were washed with PBS, and 150 µL of fresh 0.5 mg/mL MTT was added; plates were kept in the dark at 37 °C and 5 % CO_2_ for 30 min. MTT was discarded and formazan was solubilized in 200 µL DMSO (99.9 %). Absorbance was read at 540 nm (SpectraMax microplate reader; Molecular Devices, USA). Viability was expressed as % of live cells relative to control.

### 2.6. Cytotoxicity assay

Cells in 96-well plates were treated with 100 µM iron sulfate for up to 24 h. Cytotoxicity was assayed by lactate dehydrogenase (LDH) release using the CytoTox96 Non-radioactive Cytotoxicity assay kit (Promega G1780) according to the manufacturer.

### 2.7. Cell migration

Migration was assessed in Transwell inserts (Corning; 8.0-µm pore). Cells (2 × 10^4^) in serum-free medium were seeded in the upper chamber and treated with 100 µM iron sulfate for 24 h; the lower chamber contained 10 % FBS. Migrated cells were stained with 0.2 % crystal violet and counted in ImageJ.

### 2.8. Proliferation assay

Cells in 96-well plates were treated with 100 µM iron sulfate for 24 h and proliferation was measured with the BrdU Cell Proliferation ELISA kit (colorimetric; ab126556) according to the manufacturer.

### 2.9. High-resolution respirometry

Cells grown in 90-cm^2^ dishes (0.5 × 10^6^/well) and treated or not with iron were trypsinized, centrifuged (321 × g, 10 min, 21.5 °C) and resuspended in 2 mL FBS-free MEM-F12. Cells were counted (Neubauer chamber) and viability checked by Trypan blue. A volume corresponding to 2 × 10^6^ cells was added to each chamber of a high-resolution O2k respirometer (Oroboros Instruments, Austria) at 37 °C, filled to 2 mL with FBS-free MEM-F12 and equilibrated ∼10 min (stirring 750 rpm). Oxygen consumption rates were recorded in DatLab 6.0 and normalized to cell number. After basal stabilization, 500 ng/mL oligomycin was added to inhibit F_1_F_o_ ATP synthase; maximal respiration was induced by stepwise FCCP titration (100–600 nM); complex I and III contributions were probed with 100 nM rotenone and 1 µg/mL antimycin A. The six mitochondrial respiratory states were derived as described [44]: Basal; OXPHOS (basal minus oligomycin); Leak (oligomycin minus antimycin A); Spare (FCCP-induced maximum minus basal); ETS (FCCP maximum minus antimycin A); and residual oxygen consumption (ROx; after antimycin A). To assess OXPHOS coupling, two flux control ratios (FCR) were computed [45]: the leak/ETS (L/E) ratio (0, fully coupled, to 1, fully uncoupled) and the OXPHOS/ETS (P/E) ratio (0, no OXPHOS capacity, to 1, full OXPHOS capacity).

### 2.10. Mitochondrial membrane potential (Δψ_m_)

Δψ_m_ was estimated with tetramethylrhodamine ethyl ester (TMRE; Thermo Scientific). Cells on coverslips (24-well plates) treated or not with iron were rinsed twice with PBS and loaded with 60 nM TMRE in phenol-red-free MEM for 90 min (37 °C, 5 % CO_2_); 100 nM FCCP was added in the last 15 min as a depolarization control. Cells were washed in PBS containing 60 nM TMRE and mounted in 3 µL phenol-red-free MEM with 60 nM TMRE (plus 100 nM FCCP for the control). Confocal images were acquired on a Leica DMI4000B (63×/1.3 oil; excitation 549 nm, emission 574 nm; 3× digital zoom, LAS X 3.4.2). Mean grey value was quantified in Fiji [46].

### 2.11. Mitochondrial morphology

Morphology was assessed with MitoTracker Green FM (MTG; Thermo Scientific). Cells on coverslips treated or not with iron were rinsed twice with PBS and loaded with 400 nM MTG in phenol-red-free MEM for 90 min (37 °C, 5 % CO_2_). Confocal images were acquired as above (excitation 490 nm, emission 516 nm). Cell area and mitochondrial circularity was quantified in Fiji [46]; mitochondrial content was estimated from MTG integrated density per cell.

### 2.12. Total cellular oxidant levels

Intracellular oxidants were estimated with dihydroethidium (DHE; Sigma-Aldrich). Cells on coverslips treated or not with iron were washed with PBS and loaded with 30 µM DHE in phenol-red-free MEM for 15 min (37 °C, 5 % CO_2_). Confocal images were acquired on a Leica DMI4000B (63×/1.3 oil; excitation 518 nm, emission 605 nm; 3× digital zoom). Mean grey value was quantified in Fiji [46]. As detailed in the Discussion, DHE fluorescence reports bulk oxidation rather than a single oxidant species, and this caveat is considered in the interpretation.

### 2.13. Cellular hydrogen peroxide production

H_2_O_2_ production was assessed by Amplex Red oxidation. Cells treated or not with iron were trypsinized, centrifuged (321 × g, 10 min, 21.5 °C) and resuspended in 1 mL buffer (125 mM NaCl, 3 mM KCl, 2 mM CaCl_2_, 1.25 mM NaH_2_PO_4_, 2 mM MgCl_2_, 20 mM HEPES). Resorufin fluorescence from oxidation of 5 µM Amplex Red (Thermo Scientific) in the presence of 20 mg/mL horseradish peroxida se (HRP; Sigma-Aldrich) was monitored on a Cary Eclipse spectrofluorometer (Varian, USA) with stirring (excitation 530 nm, emission 590 nm); an H_2_O_2_ standard curve was run after each measurement. The well-described confounders of the Amplex Red/HRP assay (oxidation driven by NADH and reduced glutathione, and optical/redox interference by iron) are addressed in the Discussion an d are the basis for presenting the reduced-oxidant finding as a working model.

### 2.14. Sample preparation and protein digestion for proteomics

SH-SY5Y cells maintained in MEM + F-12 (1:1) with 10 % FBS and penicillin/streptomycin (10 U/mL) at 37 °C and 5 % CO_2_ were treated or not with 100 µM iron sulfate (CAS 7782-63-0; Sigma-Aldrich) for 24 h. Cells were washed with PBS and detached with trypsin/EDTA. Total lysates were prepared in RIPA buffer (R0278, Sigma-Aldrich) with protease inhibitor cocktail (P8340). Protein was quantified by BCA (23225, Pierce). Ten µg per sample were solubilized in Laemmli buffer, resolved by SDS-PAGE, stained with colloidal blue, and each lane cut into 8 bands, then into 1 × 1 mm pieces. Gel pieces were destained in 25 mM ammonium bicarbonate/50 % ACN, rinsed, shrunk in ACN, dried, and digested with trypsin (10 ng/µL in 40 mM NH_4_HCO_3_/10 % ACN; 4 °C 10 min, then 37 °C overnight). Peptides were extracted twice with H_2_O/ACN/HCOOH (47.5:47.5:5), dried and resuspended in 100 µL H_2_O/HCOOH (100:0.1).

### 2.15. nLC-MS/MS analysis

Peptide mixtures were analyzed using an Ultimate 3000 nanoLC system (Dionex, Amsterdam, The Netherlands) coupled to an Orbitrap Fusion™ Lumos™ Tribrid™ mass spectrometer (Thermo Fisher Scientific, San Jose, CA). Ten µL of digest were loaded onto a 300-µm × 5-mm C18 PepMap trap (LC Packings) at 10 µL/min and eluted onto a 75-µm × 15-cm C18 PepMap column with a 4–40 % linear gradient of solvent B over 48 min (A: 0.1 % formic acid in 5 % ACN; B: 0.1 % formic acid in 80 % ACN) at 300 nL/min. The instrument operated in positive mode (2.0-kV needle). MS scans (m/z 375–1500) were recorded at R = 120 000 (AGC 4 × 10^5^, 50 ms). Dynamic exclusion was 60 s and top speed fragmentation in HCD mode was performed over a 3 s cycle. MS/MS scans used an AGC target of 3 × 10^3^ (max fill 300 ms).

### 2.16. Database search and processing

Data were searched with SEQUEST in Proteome Discoverer 1.4 (Thermo Fisher Scientific) against the 2017.10 UniProt Human Reference Proteome Set (71,294 entries). Spectra from peptides >5000 Da or <350 Da were rejected. Precursor and fragment tolerances were 10 ppm and 0.6 Da; only b- and y-ions were used. Methionine oxidation (+16 Da) and cysteine carbamidomethylation (+57 Da) were variable modifications; up to two missed cleavages were allowed. Peptides were validated with Percolator at a 1 % false discovery rate (FDR) at the peptide level.

### 2.17. Label-free quantification and pathway analysis

Raw data were imported into Progenesis LC-MS 2.0 (Nonlinear Dynamics, UK): feature detection, alignment across all acquired LC-MS runs, integration of 2–6 charge-state ions, normalization to total protein abundance, sequence import, peptide-level ANOVA (*p*<0.05), protein abundance calculation (sum of corresponding peptides), and protein-level ANOVA (p<0.05). Only non-conflicting features and unique peptides were used, and proteins quantified with ≥ 2 peptides were retained. Four biological replicates per group were analyzed. Proteins with significantly altered abundance between groups were interrogated with Ingenuity Pathway Analysis (IPA; Qiagen) using the core analysis and upstream-regulator (“regulators”) modules.

### 2.18. qPCR and mitochondrial DNA content

Total DNA was extracted with the DNeasy Blood & Tissue kit (Qiagen) according to the manufacturer’s protocol and quantified with a NanoDrop spectrophotometer. Two nanograms of total DNA were used in multiplex qPCR reactions with TaqMan assays targeting the mitochondrial *MT-RNR1* gene (Hs02596859_g1) and the nuclear *B2M* gene (Hs06637353_s1). Mitochondrial DNA content was expressed relative to nuclear (genomic) DNA by the 2^−ΔCt^ method. Data are expressed as mean ± SEM of independent biological replicates (n = 3-9) and were analyzed as described in Section 2.20.

### 2.19. RNA extraction, cDNA synthesis and RT-qPCR

Total RNA was extracted from SH-SY5Y cells with the Monarch Total RNA Miniprep Kit (New England Biolabs, #T2010). RNA purity was verified with a NanoDrop spectrophotometer (A260/280 and A260/230 ratios > 1.8). First-strand cDNA was synthesized from 1 µg of total RNA with the SuperScript IV VILO Master Mix (Invitrogen, Cat. No. 11756050). Quantitative real-time PCR was performed on an Applied Biosystems 7500 Real-Time PCR System (Thermo Fisher Scientific) with TaqMan assays for *LARP1*(Hs00326039_m1) and the reference gene *GUSB* (Hs00939627_m1) in multiplex reactions, and relative *LARP1* expression was calculated by the 2^−ΔΔCt^ method.

### 2.20. Data and statistics

Data are presented as mean ± standard error of the mean (SEM) of n independent experiments, with n specified in each figure legend. The normality of data distribution was assessed by the D’Agostino–Pearson test, and outliers were identified by Grubbs’ test whenever a Gaussian distribution was confirmed. Normally distributed data were compared by the unpaired Student’s t-test (two groups) or by one-way ANOVA followed by Tukey’s post hoc test (three or more groups); data that deviated from normality were compared by the Mann–Whitney test. Differences were considered statistically significant at *p*<0.05. All analyses were performed in GraphPad Prism 6.00 (GraphPad Software, San Diego, CA, USA).

### 2.21. Use of AI-assisted technologies

During the preparation of this work, the authors used the large language model Claude (Anthropic; Opus 5 and Opus 5.5 models, June and September 2026) to improve the readability and language of the manuscript and to assist in formatting the reference list according to the journal style. After using this tool, the authors reviewed and edited the content as needed and took full responsibility for the content of the publication.

## 3. Results

### 3.1. Non-toxic iron overload promotes proliferation while suppressing migration

To define a non-toxic iron condition, we first measured viability across iron-sulfate concentrations (24 h). Significant losses of viability occurred only above 1 mM (EC50 ≈ 1.5 mM) (Fig. 1A). To keep cell death below 15 %, we selected 100 µM iron sulfate for all subsequent experiments. At this concentration, cellular morphology (Fig. 1B), cell area (Fig. 1C), cytotoxicity assessed by extracellular LDH release (Fig. S1A, Online Resource 1) and total protein content (Fig. S1B, Online Resource 1) were unchanged, confirming the absence of overt toxicity. ICP-MS showed an approximately four-fold increase in intracellular iron after treatment (Fig. 1D). Functionally, iron reduced migration by ∼37 % (Fig. 1E) while increasing proliferation ∼3.5-fold (Fig. 1F). Thus, under these conditions, a non-cytotoxic iron load promotes a proliferative phenotype at the expense of migratory capacity.

**Fig. 1.**
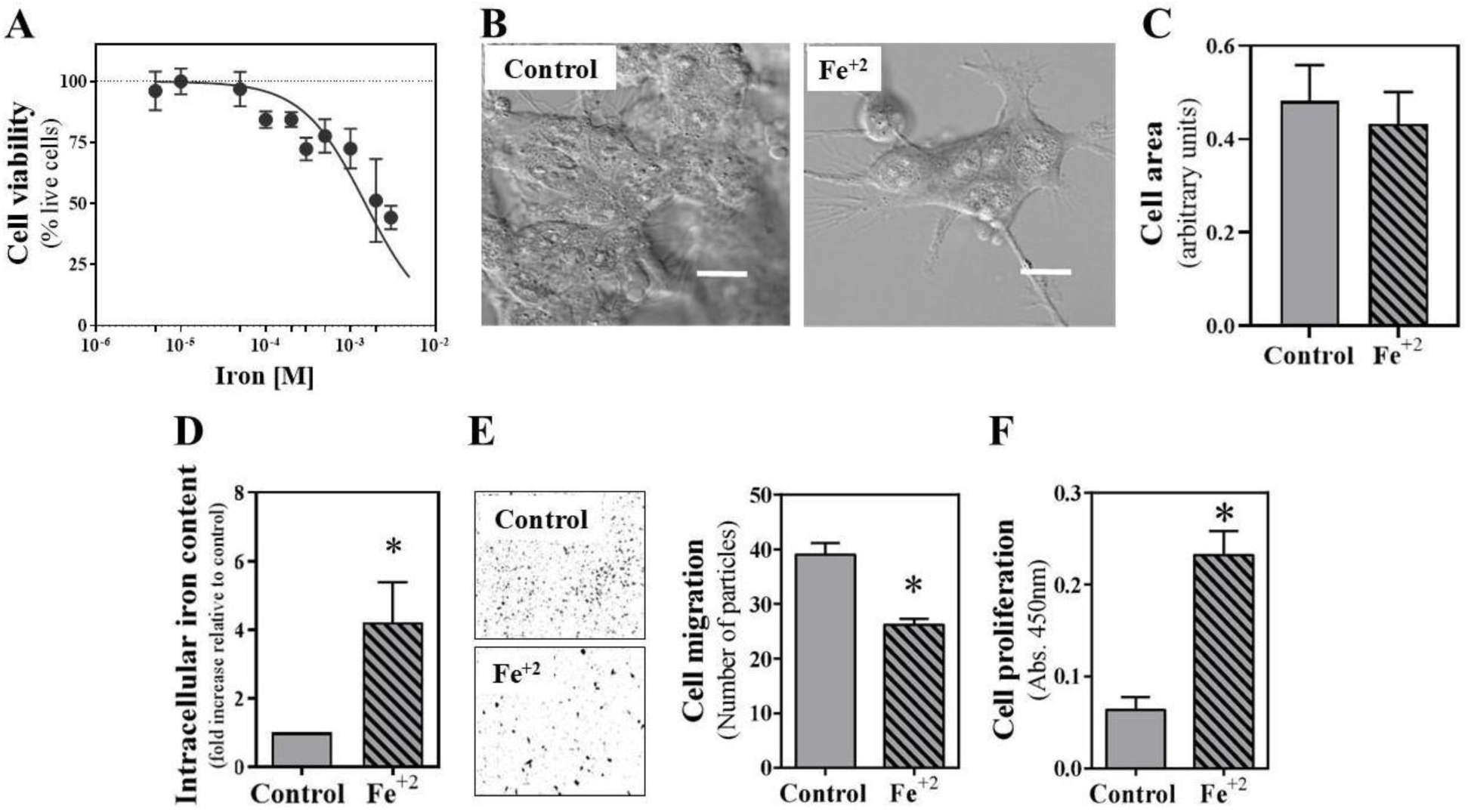
Non-toxic iron overload induces proliferation and reduces migration in neuroblastoma cells. **A.** Cell viability of neuroblastoma cells treated for 24 h with the indicated iron concentrations, assessed by MTT reduction. For **(B–F)**, cells were treated for 24 h with 100 µM iron or maintained as control. **(B)** Representative cell morphology imaged by differential interference contrast (DIC) microscopy; scale bars = 10 µm. **(C)** Cell area quantified in Fiji from confocal microscopy images. **(D)** Intracellular iron content measured by ICP-MS. **(E)** Cell migration determined by transwell assay. **(F)** Proliferation determined by BrdU incorporation assay. In all bar graphs, grey bars denote control and hatched bars denote 100 µM iron–treated cells. Data are expressed as the mean ± SEM of at least three independent experiments. \**p*<0.05 versus control (Mann–Whitney test).

### 3.2. Iron induces mitochondrial fragmentation and Δψ_m_ depolarization while preserving organelle content

Since mitochondria are a major iron store and central to energy and redox metabolism [47], we examined mitochondrial structure and function by confocal microscopy using MTG and TMRE. Iron did not change mitochondrial content, estimated either as the mtDNA/nuclear DNA ratio (Fig. 2C) or as MTG integrated density per cell (Fig. 2A, B) and per cell area (Fig. S1C, Online Resource 1). Iron produced a slight but significant increase in mitochondrial circularity, indicating a shift toward more fragmented structures (Fig. 2D), and markedly reduced Δψ_m_ (TMRE) by ∼55 % (Fig. 2A, E). Because MTG accumulation can retain some Δψ_m_ dependence, and Δψ_m_ falls by ∼55 % here, the potential-independent mtDNA/nDNA ratio provides a more robust estimate of mitochondrial content. Thus, mild iron loading remodels mitochondrial morphology and lowers Δψ_m_ without altering organelle content.

**Fig. 2.**
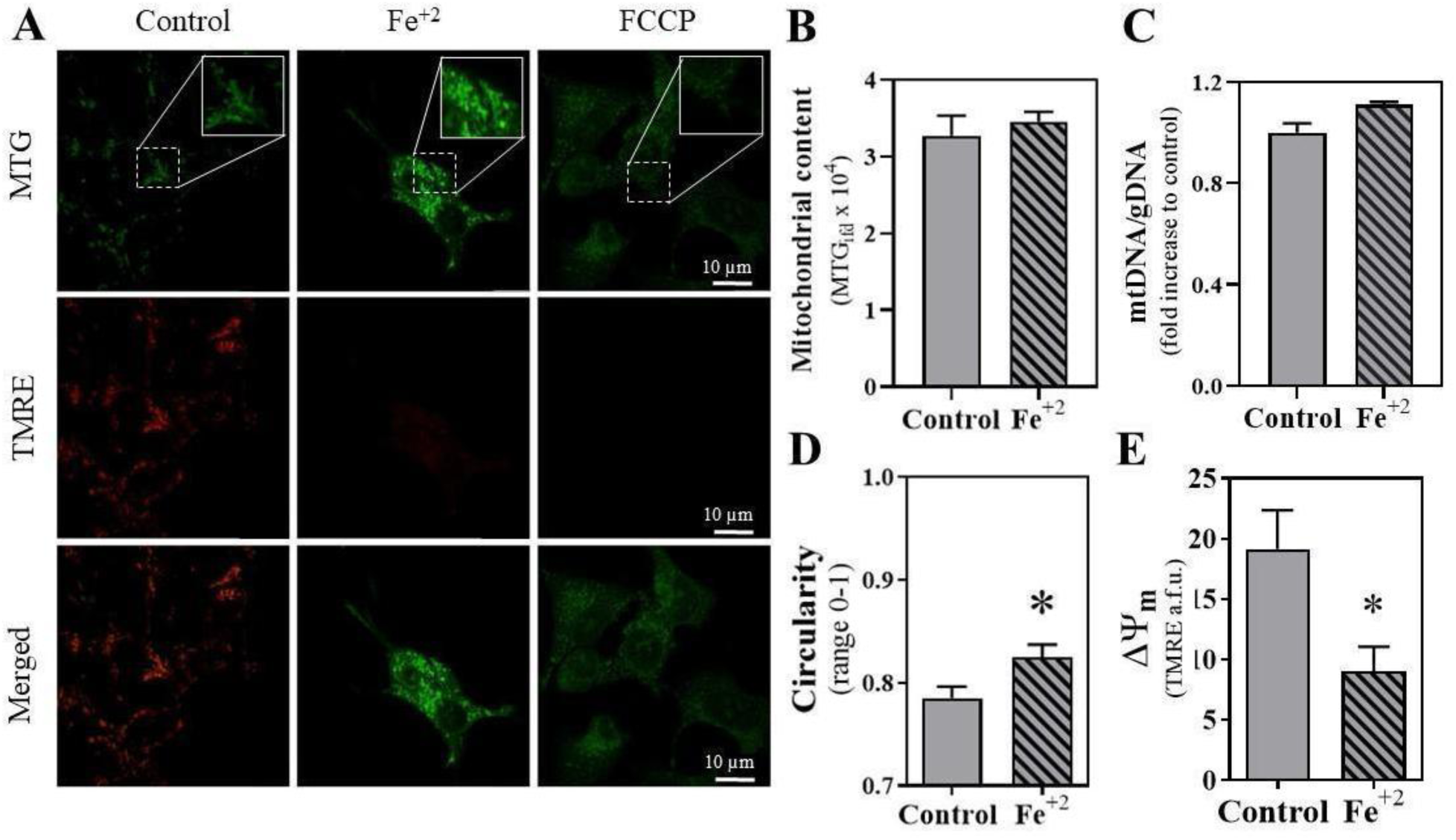
Non-toxic iron overload promotes mitochondrial fragmentation and reduces mitochondrial membrane potential in neuroblastoma cells. **A**. Neuroblastoma cells were treated for 24 h with 100 µM iron, 100 nM FCCP, or vehicle (control). **(A)** Representative confocal fluorescence microscopy images of mitochondrial morphology and membrane potential (Δψ_m_), visualized with MitoTracker Green (MTG) and TMRE, respectively. **(B, C)** Mitochondrial content, assessed as (B) the MTG integrated fluorescence density of the confocal images, quantified in Fiji, and **(C)** the mitochondrial-to-nuclear DNA ratio measured by qPCR. **D.** Mitochondrial circularity was quantified in Fiji from the confocal images of MTG-stained cells. **E.** Δψ_m_, quantified as the TMRE fluorescence intensity of the confocal images in Fiji. In all bar graphs, grey bars denote control and hatched bars denote 100 µM iron–treated cells. Data are expressed as the mean ± SEM of at least ten independent experiments. \**p*<0.05 versus control (Mann–Whitney test).

### 3.3. Iron selectively increases mitochondrial proton-leak respiration

We next used high-resolution respirometry in intact cells, resolving six mitochondrial states (Fig. 3A) from representative oxygen-consumption traces of control and iron-treated cells (Fig. 3B). Across all six states, iron had minimal impact on respiration overall (Fig. 4A–F), with a single, significant and specific increase in proton-leak respiration (Fig. 4C), consistent with the Δψ_m_ depolarization in Fig. 2E. Because mitochondrial content was unchanged (Fig. 2B, C), we also normalized respiratory rates to the mtDNA/nuclear DNA ratio; again, only proton-leak respiration was significantly increased (Fig. 5C). Mild iron overload therefore promotes proton-leak respiration independently of mitochondrial content.

**Fig. 3.**
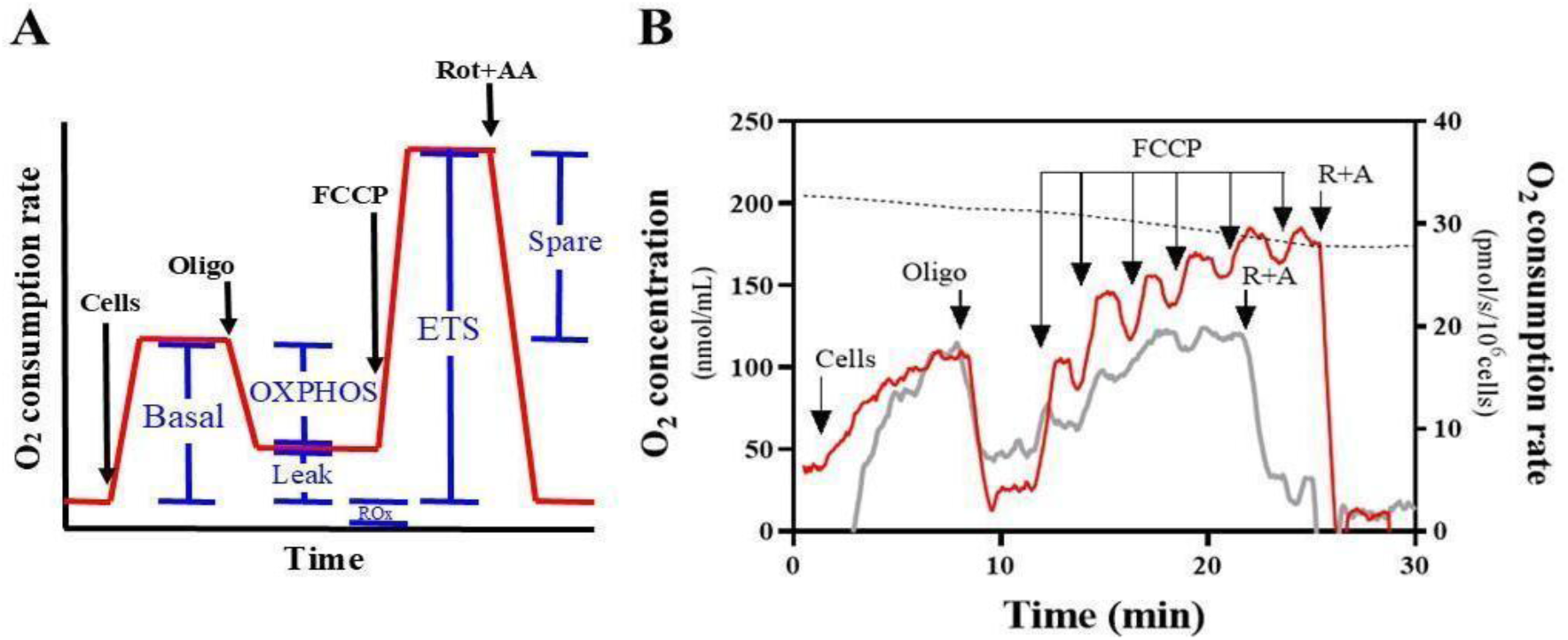
Assessment of oxygen consumption of intact neuroblastoma cells. A. Schematic of a typical oxygen-consumption-rate trace (red line) of intact cells obtained by high-resolution respirometry. Arrows mark the additions of OXPHOS modulators. The six mitochondrial respiratory states (basal, OXPHOS, leak, spare, ETS and ROx) were derived as described in Methods and are labelled in blue. **B.** Representative oxygen-consumption-rate traces of intact control (red line) and 100 µM iron-treated (grey line) neuroblastoma cells, measured with 2 × 10⁶ cells in 2 mL of FBS-free MEM-F12 in the chamber of an O2k respirometer at 37°C. Arrows mark the additions of OXPHOS modulators, at the following final concentrations: oligomycin (oligo), 500 ng/mL; FCCP, 100 nM per injection; rotenone (rot), 100 nM; antimycin A (AA), 1 µg/mL. The dotted line shows the oxygen tension in the chamber (left y-axis).

**Fig. 4.**
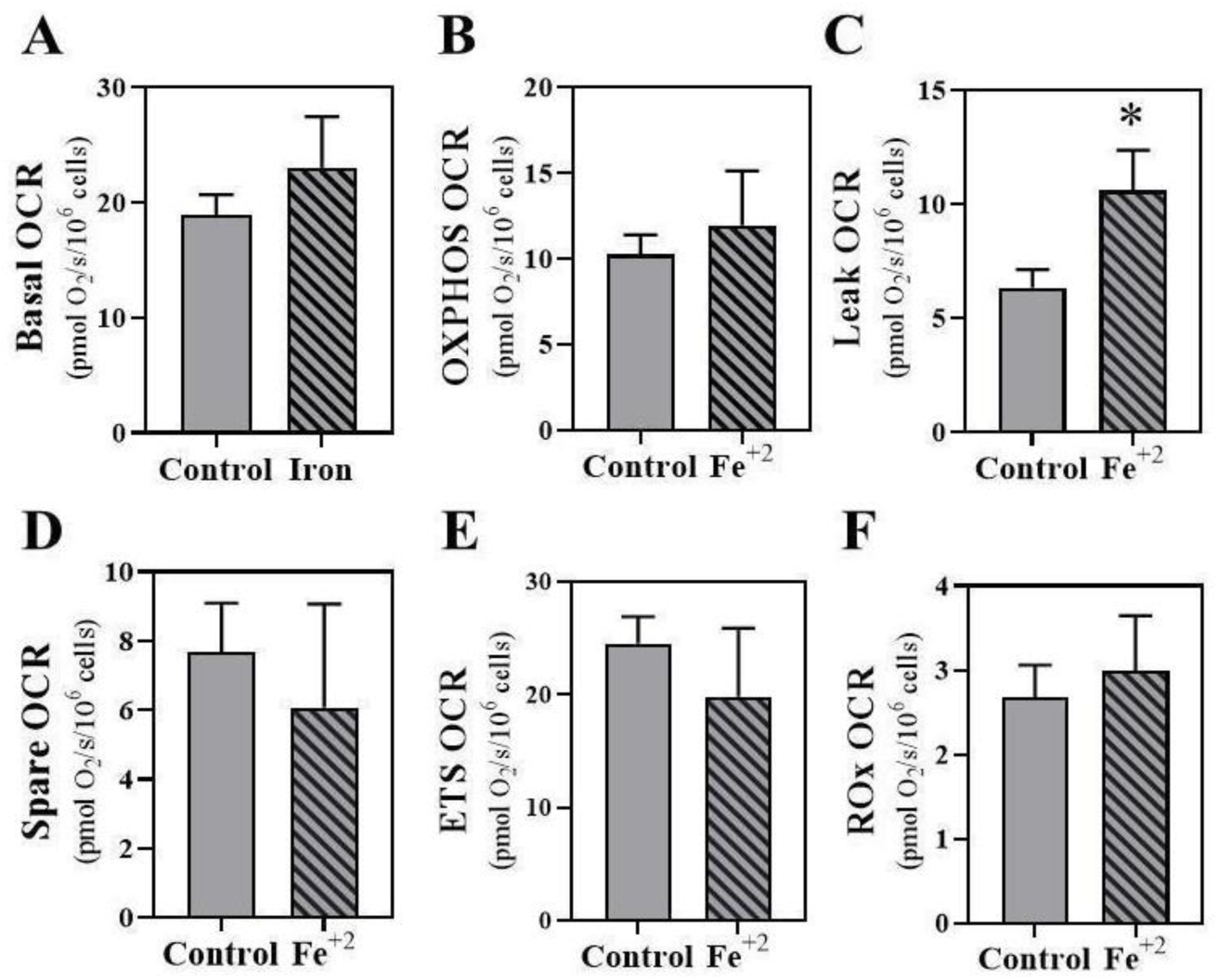
Effect of iron on mitochondrial respiratory states. Oxygen consumption rates (OCR; O₂ flow per cell, pmol·s⁻¹·10⁻⁶ cells) in the basal (A), OXPHOS (B), leak (C), spare (D), ETS (E) and ROx (F) states were determined in control and 100 µM iron-treated neuroblastoma cells. Grey bars denote control and hatched bars denote 100 µM iron-treated cells. Data are presented as the mean ± SEM of at least three independent experiments. \**p*<0.05 versus control (Mann–Whitney test).

**Fig. 5.**
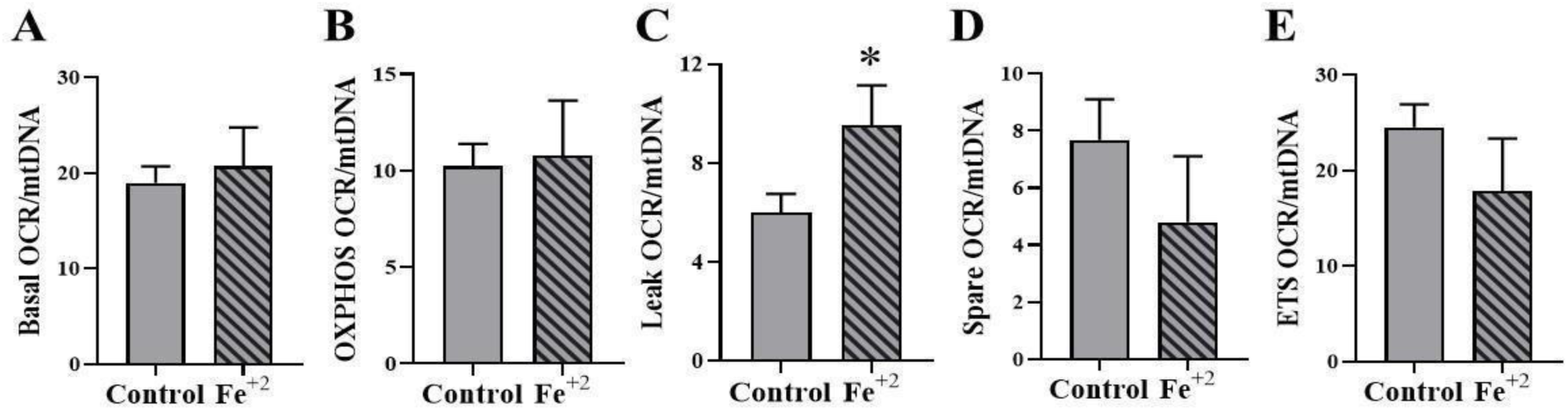
Iron-promoted increase in proton-leak respiration is independent of mitochondrial content. Mitochondrial respiratory rates in the basal, OXPHOS, leak, spare and ETS states were normalized to the mtDNA/gDNA ratio as a read-out of mitochondrial content. In all bar graphs, grey bars denote control and hatched bars denote 100 µM iron-treated cells. Data are presented as the mean ± SEM of at least five independent experiments. \**p*<0.05 versus control (Mann–Whitney test).

### 3.4. OXPHOS coupling efficiency is preserved upon iron treatment

To test whether the increased leak compromised coupling between respiration and ATP synthesis, we calculated the leak/ETS (L/E) and OXPHOS/ETS (P/E) flux control ratios in intact cells (Fig. 6). In control cells, the contributions of leak and OXPHOS to maximal ETS respiration were relatively high (L/E ≈ 0.25; P/E ≈ 0.40). Iron significantly increased the L/E ratio (Fig. 6A), reinforcing the increase in proton-leak respiration (Figs. 4C and 5C), whereas the P/E ratio rose only slightly and non-significantly (Fig. 6B). Hence, OXPHOS coupling efficiency is preserved despite the higher proton leak. A 55 % fall in Δψm together with “minimal impact” on basal/OXPHOS oxygen flux is best explained by a compensatory increase in electron-transport flux that maintains net phosphorylation efficiency at a lower Δψ_m_.

**Fig. 6.**
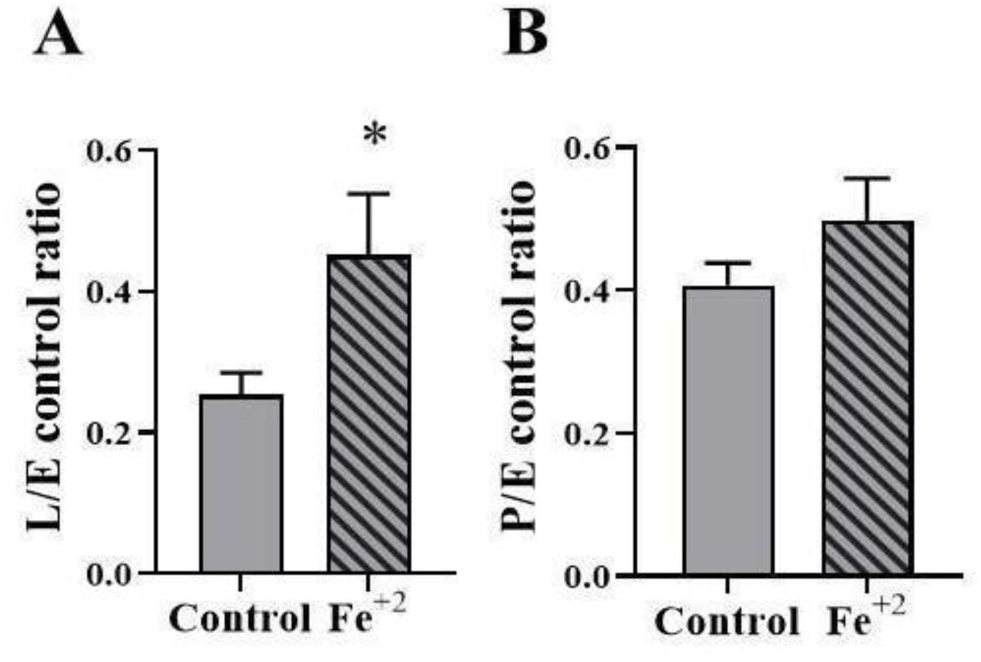
Effect of iron on flux control ratios. (A) Leak/ETS (L/E) and (B) OXPHOS/ETS (P/E) flux control ratios were determined in control and 100 µM iron-treated neuroblastoma cells. Grey bars denote control and hatched bars denote 100 µM iron-treated cells. Data are presented as the mean ± SEM of at least three independent experiments. \**p*<0.05 versus control (Mann–Whitney test).

### 3.5. Iron treatment reduces cellular oxidant production

Mitochondrial oxidant production is strongly governed by Δψ_m_ [48–49]. Given the higher proton leak and lower Δψ_m_ induced by mild iron exposure, we asked whether iron altered cellular oxidant levels. Total DHE fluorescence decreased upon iron treatment (Fig. 7A), with quantification indicating a ∼55 % reduction in oxidant signal (Fig. 7B). Consistently, cellular H_2_O_2_ production, followed by Amplex Red oxidation, was strongly reduced (Fig. S1D, Online Resource 1). Because both probes are subject to known confounders in an iron-loading context (see Discussion), we interpret these concordant decreases as a robust but probe-based working model of reduced oxidant output rather than as a fully established quantitative measurement of a single oxidant species.

**Fig. 7.**
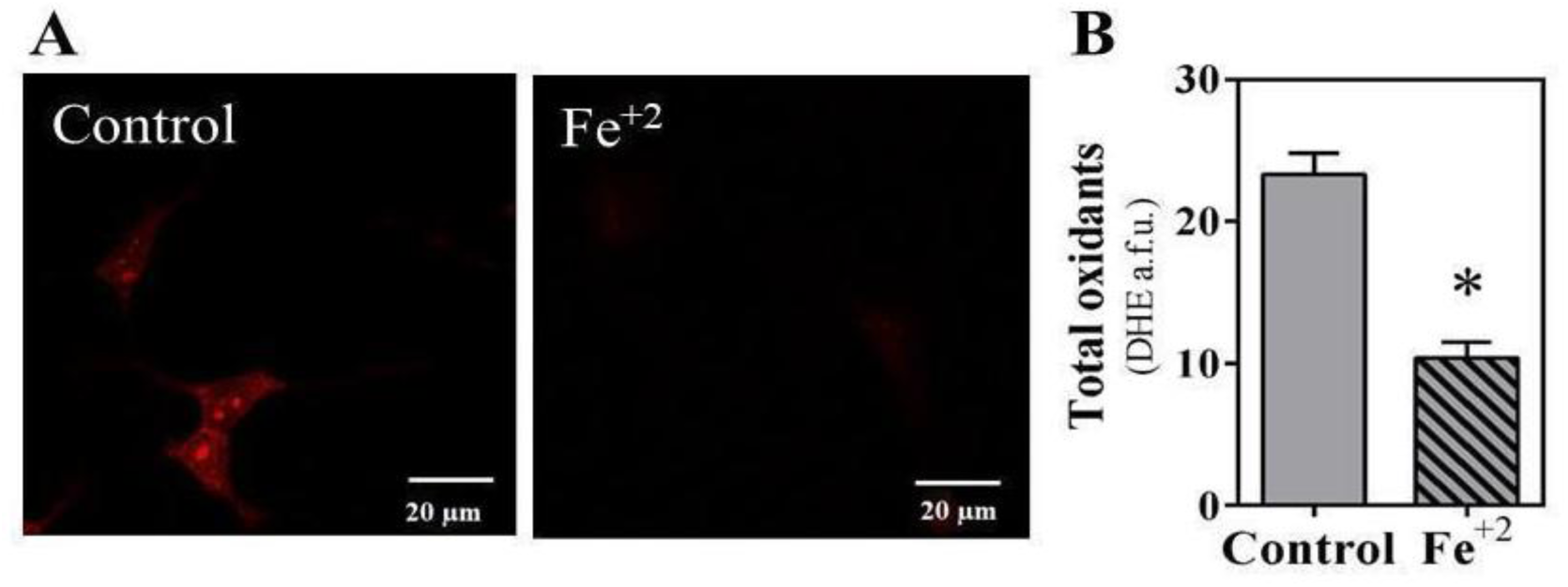
Iron exposure reduces total cellular oxidant production in neuroblastoma cells. A. Representative confocal fluorescence microscopy images of total oxidant levels in control and 100 µM iron-treated neuroblastoma cells, visualized with dihydroethidium (DHE). **B.** Total oxidant levels, quantified as DHE fluorescence intensity from confocal images using Fiji and expressed in arbitrary fluorescence units (a.f.u.). Grey bars, control; hatched bars, 100 µM iron-treated cells. Data are presented as the mean ± SEM of at least five independent experiments. \**p*<0.01 versus control (Mann–Whitney test).

### 3.6. Iron remodels neuroblastoma proteome toward reduced translation

Label-free quantitative mass spectrometry (24 h; three biological replicates per group) identified 129 proteins with significantly altered abundance (p<0.05): 25 increased and 104 decreased (Fig. 8). The increased proteins were dominated by core histones (e.g., H3, H4, H2B and H2A species) and by both ferritin light and heavy chains, together with the mitochondrial ATP synthase subunit ε-like protein (ATP5EP2). The decreased proteins were enriched in RNA-splicing factors, translation machinery and lipid-metabolism enzymes, and notably included transferrin receptor protein 1 (TfR1), consistent with a cell adapting to higher iron availability. Among ribosomal proteins, the consistent direction of change was downward (for example, the 60S subunits L37 and L39 and the 40S subunits S26 and S30 were reduced; Fig. 8). IPA of the 129 altered proteins identified protein translation and RNA processing as the most significantly affected pathways, with the most enriched terms (rRNA processing, eukaryotic translation initiation/elongation/termination, nonsense-mediated decay and EIF2 signaling) showing negative activation z-scores, i.e., a predicted decrease (Fig. 9). Upstream-regulator analysis (Table 1) predicted activation of LARP1 (z ≈ +4.2) and RICTOR (z ≈ +3) and inhibition of MLXIPL (z ≈ −4), MYCN, MYC and related regulators, with ribosomal proteins recurring as their shared target molecules. We emphasize that these upstream regulators are computational predictions.

**Fig. 8.**
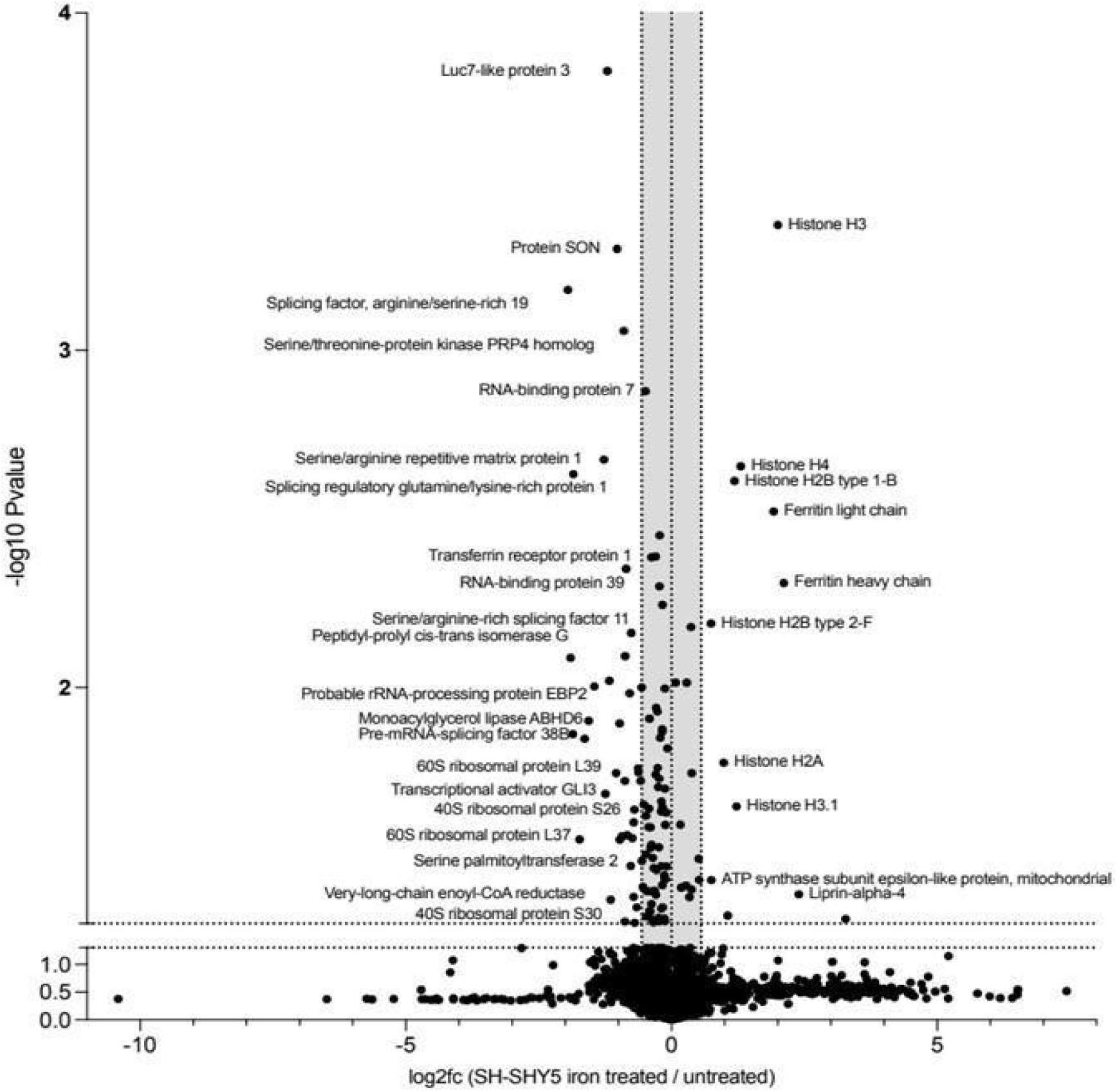
Changes in neuroblastoma proteome promoted by iron treatment. Label free unbiased differential proteomic study of neuroblastoma cells exposed to 100 µM iron for 24 h as compared to control cells. The repressed proteins are shown on the lef t side of the Volcano Plot and the over-represented proteins on the right. The X axis gives the quantitative change expressed by log2 of the fold change (iron vs, control). The Y axis provides the significance of the changes expressed as the -log10 of the adjusted *p* value (n=4).

**Fig. 9.**
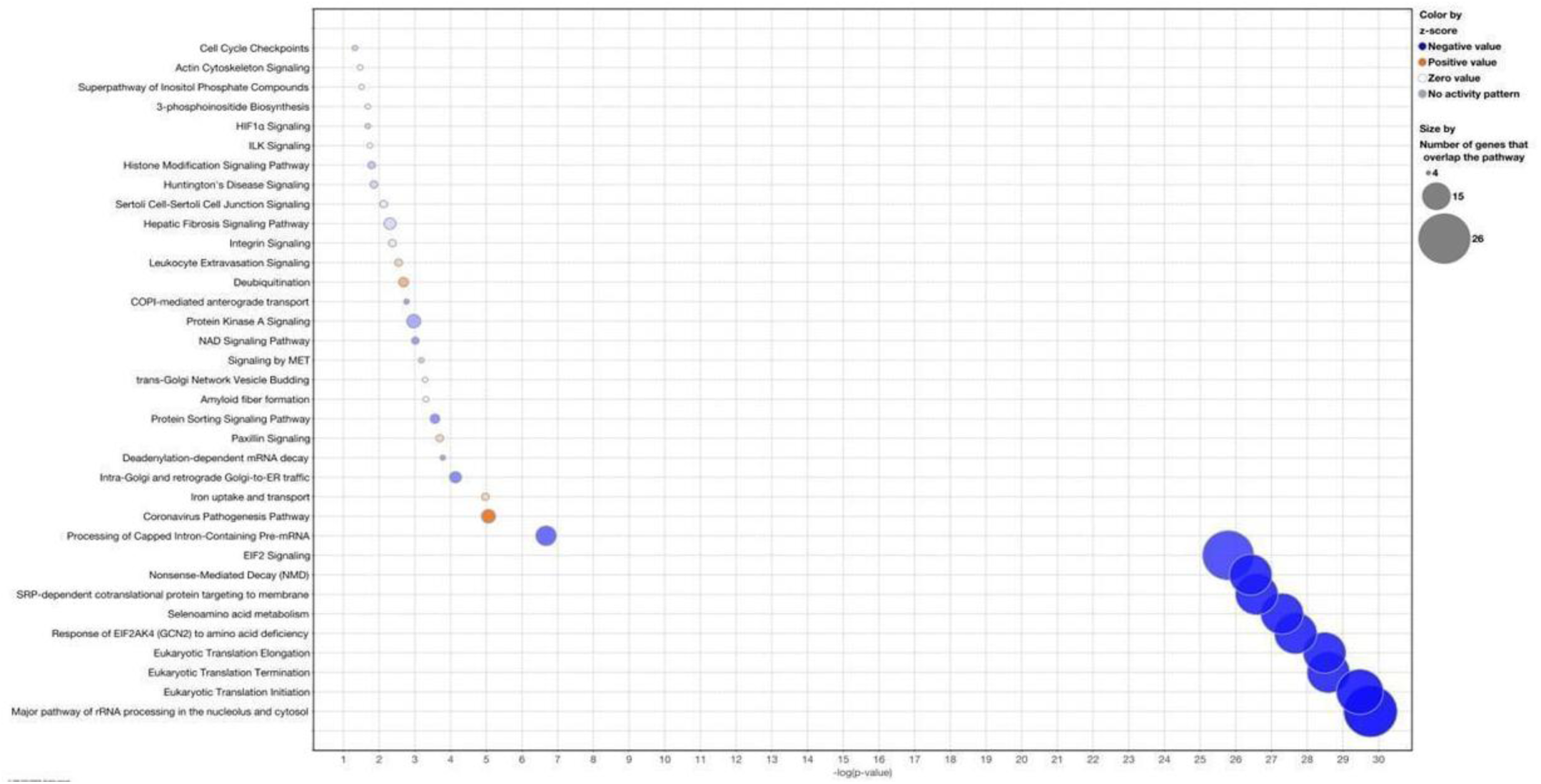
The pathway analysis is shown as a bubble volcano plot (significant pathways with -logAdjPvalue>1.3 are shown. The pathways with blue dots are inhibited while pathways with orange dots are activated. The number of proteins detected for each pathway is represented by the diameter of each dot. Activation or inhibition was determined using the Z-score calculated by IPA Qiagen.

**Table 1.**
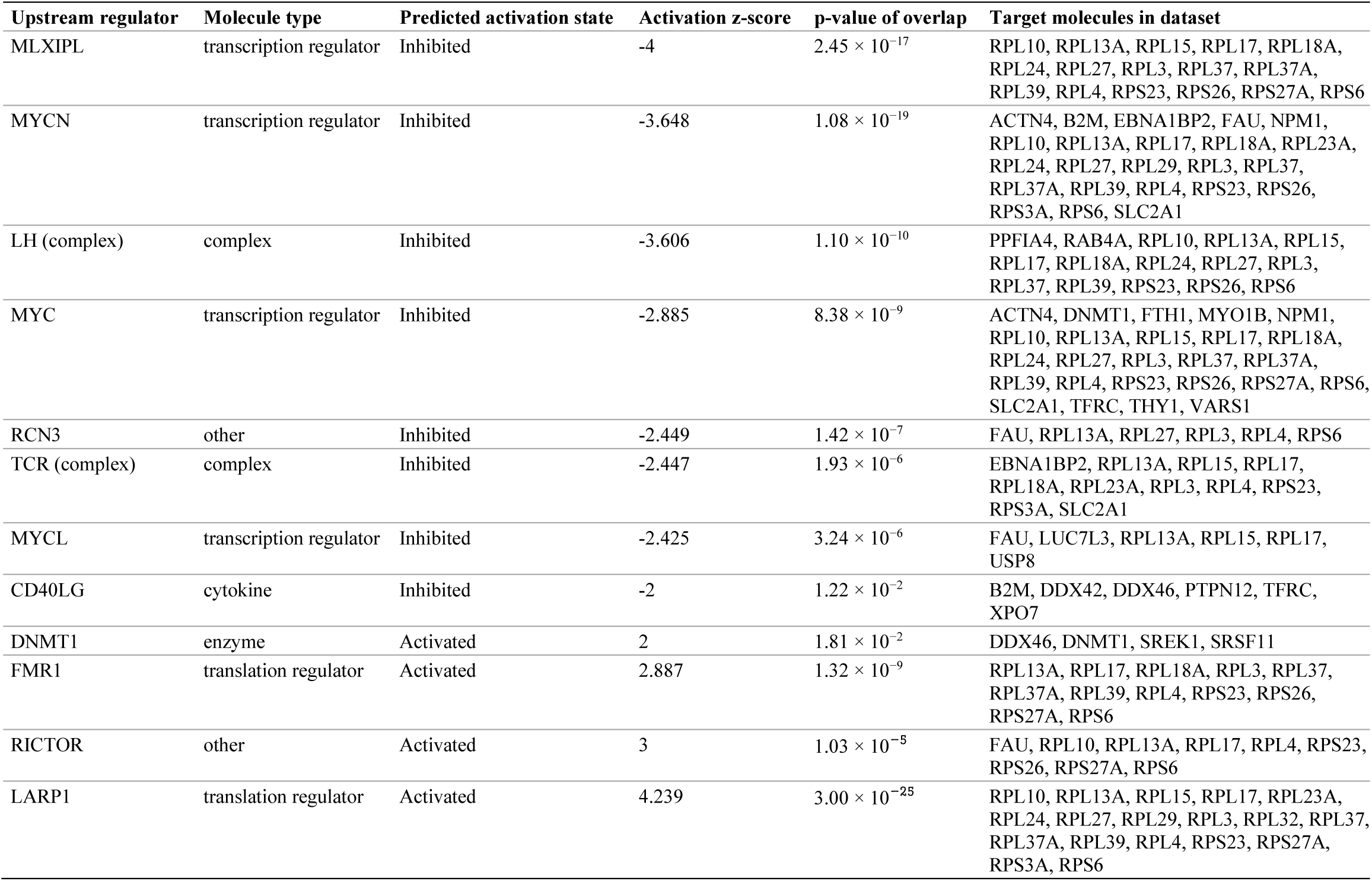
Upstream regulators predicted by Ingenuity Pathway Analysis from the proteome of iron-treated neuroblastoma cells. Regulators are ranked by activation z-score, with their predicted target proteins and overlap p-value; only regulators with |z| ≥ 2 and p<0.05 are shown.

As a first partial test of the translational arm, LARP1 transcript levels were significantly increased upon iron treatment by RT-qPCR (Fig. S1E, Online Resource 1), even though LARP1 protein was not significantly altered in the proteome.

## 4. Discussion

Here, we investigated how sub-lethal iron exposure affects neuroblastoma cells, assessing cellular, metabolic, and molecular parameters to clarify its potential consequences for tumor progression and migration. Our central, data-constrained finding is a coherent bioenergetic axis: a mild, sub-cytotoxic iron load selectively increases mitochondrial proton-leak respiration and lowers Δψ_m_, accompanied by a marked reduction in cellular oxidant signals, while OXPHOS coupling efficiency and mitochondrial content are preserved. In parallel, mild iron exposure promotes neuroblastoma proliferation while reducing migration, a phenotype consistent with the well-documented dependence of these cells on iron for viability and proliferation [32–35, 37, 42]. Together, these observations position iron, a micronutrient cue, as a cell-autonomous driver of neuroblastoma reprogramming at the level of three cancer hallmarks [12]: energy metabolism, cell proliferation and cell migration. Notably, comparable changes have been reported in iron-overloaded rats, whose liver mitochondria showed reduced respiratory control ratios and decreased lipid peroxidation while preserving ADP/O ratios and ATP levels [40]. Collectively, these findings support the view that, rather than acting solely as a terminal source of toxic free radicals, labile iron can drive functional mitochondrial changes that lower oxidant production, a preventive antioxidant response that offsets the pro-oxidant burden of this transition metal within the cell.

The leak–Δψ_m_–oxidant relationship is the mechanism our data most directly constrain. Mitochondria are not only the main intracellular store of iron [47] but also the principal site of cellular oxidant production, and the rate of superoxide and H_2_O_2_ generation by the electron transport system is steeply dependent on Δψ_m_. A high proton-motive force favors single-electron reduction of oxygen at complexes I and III, whereas lowering Δψ_m_, including by increasing proton conductance, sharply reduces electron leak to oxygen; this steep dependence and the antioxidant effect of mild uncoupling were established directly in isolated mitochondria, where protonophores, ADP and physiological uncouplers such as free fatty acids each suppressed H_2_O_2_ output [48, 49]. Crucially, the same logic has been demonstrated specifically for iron: in liver mitochondria from acutely iron-loaded rats, iron triggered a proton leak that lowered the respiratory control ratio yet left phosphorylation efficiency and ATP levels intact, leading the authors to propose mitochondrial uncoupling as a protective, redox-homeostatic response to iron [40].

Iron handling by mitochondria provides the context for these observations. Mitochondria are a major site of cellular iron storage and the obligatory site of iron–sulfur cluster (ISC) and heme biosynthesis [19, 47, 50–52], and iron import across the inner membrane is mediated by mitoferrins and driven by Δψ_m_ [53–56]. Iron availability therefore scales mitochondrial function in both directions. At physiological levels, iron supports mitochondrial protein synthesis and organelle biogenesis [57–59], whereas iron deprivation impairs mitochondrial translation and respiration and reshapes energy metabolism, notably in cancer cells [60–63]. At the other extreme, iron overload causes tissue injury and molecular oxidation [64–66], and oxidant production, mitochondrial fragmentation, redox imbalance and OXPHOS impairment have been reported in isolated mitochondria from several tissues [67–72]. In neuroblastoma cell lines specifically, iron overload raises mitochondrial iron and oxidant production, decreases AMPK activity and promotes α-synuclein aggregation [35, 36, 42].

Between these extremes, cells and mitochondria tolerate moderately elevated iron to a considerable extent, even when redox balance is perturbed [64, 72, 73], and the adaptive uncoupling of iron-loaded liver mitochondria discussed above [40] is one expression of this tolerance. Conversely, iron depletion by chelation triggers mitophagy and lowers mitochondrial content [41], a response opposite to the preserved mitochondrial content measured here (Fig. 2B, C). Our data therefore place mild iron loading within a window of mitochondrial metabolic flexibility in which bioenergetic remodeling limits redox damage and preserves viability, clearly distinct from the injurious regime of frank overload.

In this framework, the iron-induced increase in proton conductance produces a biophysically sufficient antioxidant effect: by dissipating a portion of Δψ_m_, it reduces the thermodynamic driving force for single-electron reduction of oxygen, lowering superoxide and H_2_O_2_ generation without any change in antioxidant-enzyme expression. This model reconciles the paradox of reduced ROS under iron loading through a purely bioenergetic mechanism: enhanced proton leak lowers Δψ_m_ to a range in which electron leak is minimized, a strategy that is energetically favorable because it preserves OXPHOS coupling efficiency while eliminating a potentially damaging oxidant signal. It therefore represents a fundamentally different antioxidant logic from enzyme-mediated ROS scavenging, operating continuously and proportionally through membrane biophysics rather than through regulated gene expression. In this framework, the phenotype generated by our treatment paradoxically poses a pro-oxidant stressor (iron) as a generator of a preventive antioxidant response.

The structural remodeling we observe in neuroblastoma mitochondria is consistent with a Δψ_m_-driven, fission-independent route [74, 75]. Iron increased mitochondrial circularity and depolarized Δψ_m_ without changing total mitochondrial content. We note that loss of Δψ_m_ is itself sufficient to bias the network toward fragmentation, because inner-membrane fusion is strictly Δψ_m_-dependent and depolarization promotes proteolytic inactivation of the fusion machinery (mitofusins and OPA1); the morphological change therefore need not invoke transcriptional or translational upregulation of canonical fission proteins. The preservation of mitochondrial content and coupling, alongside iron’s established role as an obligate cofactor for ISC and heme biosynthesis [47, 50–52], is in keeping with iron supporting, rather than degrading, respiratory competence under mild loading. Likewise, the upregulation of ferritin (light and heavy chains) and the downregulation of transferrin receptor 1 in our proteome are the expected hallmarks of a cell buffering an expanded labile iron pool, complementing the biophysical, leak-dependent limitation of oxidant production.

The mechanism underlying the iron-induced increase in proton leak and fall in Δψ_m_ remains to be established, and several non-exclusive possibilities deserve consideration. First, mitochondrial iron import is itself electrogenic: Fe^2+^ crosses the inner membrane through mitoferrin-1/2 in a Δψ_m_-dependent manner [53–56], and the mitochondrial calcium uniporter provides an additional entry route under iron overload [76]. Sustained cation influx would dissipate part of Δψ_m_ and would be recorded as oligomycin-insensitive (leak) respiration. However, at the modest (∼four-fold) rise in cellular iron reached here, the steady-state charge flux associated with iron uptake is expected to be small relative to the proton current generated by the ETS, making it unlikely that this mechanism alone accounts for a ∼55 % fall in Δψ_m_; quantifying mitochondrial iron and testing whether mitoferrin inhibition prevents the depolarization would resolve this point. Second, iron may increase inducible proton conductance: lipid-peroxidation products such as 4-hydroxynonenal activate uncoupling proteins and the adenine nucleotide translocase [77], and free fatty acids act as natural protonophores [49]. This route implies an early, iron-driven oxidative signal that is subsequently self-limited by the resulting depolarization, which is compatible with, but not demonstrated by, our steady-state measurements at 24 h. Third, the induction of ferritin light and heavy chains and the repression of TfR1 (Fig. 8) show that the iron-regulatory protein/iron-responsive element system was engaged [78]. Ferritin sequesters iron as a ferric mineral core of up to ∼4,500 atoms per 24-mer [79] and thereby limits the labile iron available for Fenton chemistry [80]. The alternative possibility that ferritin buffering becomes saturated, diverting excess iron into mitochondria, is not supported by our data: a four-fold increase in total cellular iron is modest relative to ferritin storage capacity, and the concomitant downregulation of TfR1 indicates that cells were actively restricting further iron uptake. Importantly, ferritin induction can explain a reduced iron-catalyzed formation of secondary oxidants, but not by itself the lower superoxide/H_2_O_2_ signals reported by DHE and Amplex Red, which are better explained by the lower Δψ_m_ [48, 49]. Ferritin-mediated iron sequestration and leak-dependent depolarization are therefore best viewed as complementary rather than alternative antioxidant arms.

At the proteome level, the reproducible signal is a coordinated reduction of ribosomal proteins and translation/RNA-processing components, with concurrent histone upregulation. The upstream regulators predicted by IPA to be activated (LARP1 and RICTOR) or inhibited (MYC, MYCN and MLXIPL) are mechanistically attractive: (i) LARP1 is a translational regulator of 5′TOP mRNAs encoding ribosomal proteins [81, 82]; (ii) RICTOR is the defining subunit of mTORC2 [83], a complex that drives the anabolic metabolism required to sustain proliferation (Fig. 1F); and (iii) the MYC/ChREBP transcriptional programs [84] converge on ribosome biogenesis and anabolic metabolism, with ribosomal proteins recurring as their shared target molecules (Table 1). We stress, however, that these are predictions inferred from the direction and magnitude of downstream changes, not direct measurements of pathway activity. Consistent with this caveat, the increase in LARP1 transcript by RT-qPCR (Fig. S1E, Online Resource 1) is supportive but not confirmatory, and gain/loss-of-function and phospho-proteomic experiments would be required to test the mTORC2/LARP1 axis directly.

The proteomic and functional phenotypes can be integrated, at least provisionally, within this bioenergetic framework. Protein synthesis is among the largest ATP sinks in mammalian cells [85], so a coordinated reduction of ribosomal proteins and translation machinery would lower the ATP demand for translation; that OXPHOS-linked respiration was nonetheless maintained suggests a redistribution of ATP turnover among cellular processes rather than a global decline in energy demand. The apparent paradox of increased proliferation alongside reduced translational capacity may be partly resolved by the nature of the readouts: BrdU incorporation reports DNA synthesis in S phase, and replication-dependent histone synthesis is tightly coupled to DNA replication [86], consistent with the histone upregulation detected here. Iron is also an essential cofactor for ribonucleotide reductase and DNA synthesis, and iron chelation arrests neuroblastoma cells in G1 [27, 34]; iron supplementation would be expected to relieve this constraint. Regarding migration, mitochondrial superoxide promotes tumor cell migration and metastasis, an effect prevented by mitochondria-targeted antioxidants [87]; the marked reduction in oxidant signals observed here therefore provides a plausible, testable link to the ∼37 % decrease in migration. Because these connections rest on concordance rather than intervention, experiments that restore Δψ_m_ or oxidant levels in iron-treated cells will be required to establish causality.

Several limitations frame our conclusions. (i) This is a single-cell-line, intrinsic-metabolism study in the non-MYCN-amplified SH-SY5Y line, at one iron concentration and a single 24-h timepoint, with no microenvironmental or *in vivo* component; generalizability — particularly to MYCN-amplified lines, three-dimensional models and in vivo settings — remains to be established. (ii) The reduced-oxidant finding rests on two probe-based, whole-cell assays, each with known caveats, and we therefore present it as a working model rather than an established quantity. DHE reports bulk oxidation and is not specific for a single oxidant; ideally the superoxide-specific 2-hydroxyethidium product would be resolved [88, 89]. The Amplex Red/HRP assay is confounded by NADH- and glutathione-driven oxidation and, critically in an iron-loading context, by iron’s own optical and redox interference [90, 91]. Specific or mitochondrially targeted probes, SOD/catalase-controlled Amplex Red measurements, and a causal test (re-polarizing the membrane or blocking the leak to restore oxidant output) would convert this model into a firm conclusion. (iii) Proteomic significance was set at *p*<0.05 without multiple-testing correction; because the IPA analysis is built on this list, its upstream-regulator predictions inherit that risk and require an FDR/q-value threshold and direct experimental validation.

Taken together, the data support a parsimonious model in which mild, non-toxic iron acts as a hormetic cue: a selective increase in proton leak lowers Δψ_m_ into a range that biophysically suppresses oxidant production, electron-transport flux compensates so that OXPHOS coupling and mitochondrial content are preserved, ferritin/TfR1 remodeling buffers the labile iron pool, and the proteome shifts toward reduced translation. This regime is distinct from frank iron overload (which can trigger ferroptosis) [92] and from iron chelation (which depletes iron-dependent processes and induces apoptosis) [38, 39]. Therapeutically, the leak-dependent, biophysical antioxidant mechanism cannot be targeted by inhibiting antioxidant enzymes; instead, interventions that re-polarize the inner membrane or block the leak conductance might re-expose iron-adapted cells to oxidant-mediated death — a prediction that follows from, but is not proven by, the present data.

## 5. Conclusion

We show that a mild, non-toxic iron load promotes neuroblastoma cell proliferation while reducing migration, in association with mitochondrial fragmentation and Δψ_m_ depolarization, a selective increase in proton-leak respiration with preserved OXPHOS coupling and mitochondrial content, and a strong, leak-compatible reduction in cellular oxidant signals. The proteome shifts toward reduced ribosomal-protein abundance and translation, with LARP1/RICTOR activation and MYC/MLXIPL inhibition predicted as upstream regulators. The data most firmly support a biophysical, proton-leak-dependent limitation of oxidant production as a cell-autonomous adaptive response to a micronutrient (iron) cue and identify this leak conductance as a candidate vulnerability worth testing for altering neuroblastoma progression.

## Supporting information

Table S1 Quantitative proteomic dataset

## Statements and declarations

### Ethics approval

This study did not involve human participants, human-derived samples requiring consent, or animals. It used the established human neuroblastoma cell line SH-SY5Y obtained from the Rio de Janeiro Cell Bank (BCRJ); accordingly, ethics-committee approval was not required.

### Consent to participate / Consent to publish

Not applicable.

### Data availability

The mass spectrometry proteomics data have been deposited to the ProteomeXchange Consortium via the PRIDE [93] partner repository with the dataset identifier PXD085256. All other data supporting the findings of this study are available from the corresponding author upon reasonable request.

### Author contributions

A.P.M.M., N.D.A., S.C., B.F.B., A.S.F., F.D.R., R.E.S., D.B.C., R.R. and M.F.O. conceived and designed the experiments. A.P.M.M., N.D.A., S.C., B.F.B. and A.S.F. performed the experiments. A.P.M.M., N.D.A., S.C., B.F.B., A.S.F., F.D.R., R.E.S., R.R. and M.F.O. analyzed the data. N.D.A., S.C., C.T., D.B.C., R.E.S., R.R. and M.F.O. contributed reagents/materials/analysis tools. A.P.M.M., N.D.A., R.R. and M.F.O. wrote the paper. All authors read and approved of the final manuscript.

## Acknowledgments

We thank Dr. Fernando de Almeida (Centro Nacional de Biologia Estrutural e Bioimagem, CENABIO) for assistance with confocal imaging and Dr. Mariana F. Rodrigues for contributions to molecular biology experiments.

## Abbreviations

ALK: anaplastic lymphoma kinase
ANT: adenine nucleotide translocase
ATRX: alpha thalassemia/mental retardation X-linked
BrdU: 5-bromo-2′-deoxyuridine
DHE: dihydroethidium
ETS: electron transport system
FBS: fetal bovine serum
FCCP: carbonyl cyanide p-(trifluoromethoxy)phenylhydrazone
FCR: flux control ratio
ICP-MS: inductively coupled plasma mass spectrometry
IPA: Ingenuity Pathway Analysis
ISC: iron–sulfur cluster
KED: kinetic energy discrimination
L/E: leak/ETS coupling control ratio
LDH: lactate dehydrogenase
MTG: MitoTracker Green FM
mtDNA: mitochondrial DNA
OXPHOS: oxidative phosphorylation
P/E: OXPHOS/ETS coupling control ratio
RCR: respiratory control ratio
ROS: reactive oxygen species
ROX: residual oxygen consumption
TERT: telomerase reverse transcriptase
TfR1: transferrin receptor 1
TMRE: tetramethylrhodamine ethyl ester
Δψ_m_: mitochondrial inner membrane potential.

## Supplementary information

Supplementary Fig. S1 is provided as Online Resource 1.

Supplementary Table S1. is provided as Online Resource 2.

