## Supplemental Figure S1 for "Iron exposure reprograms redox homeostasis through mitochondrial depolarization in neuroblastoma"

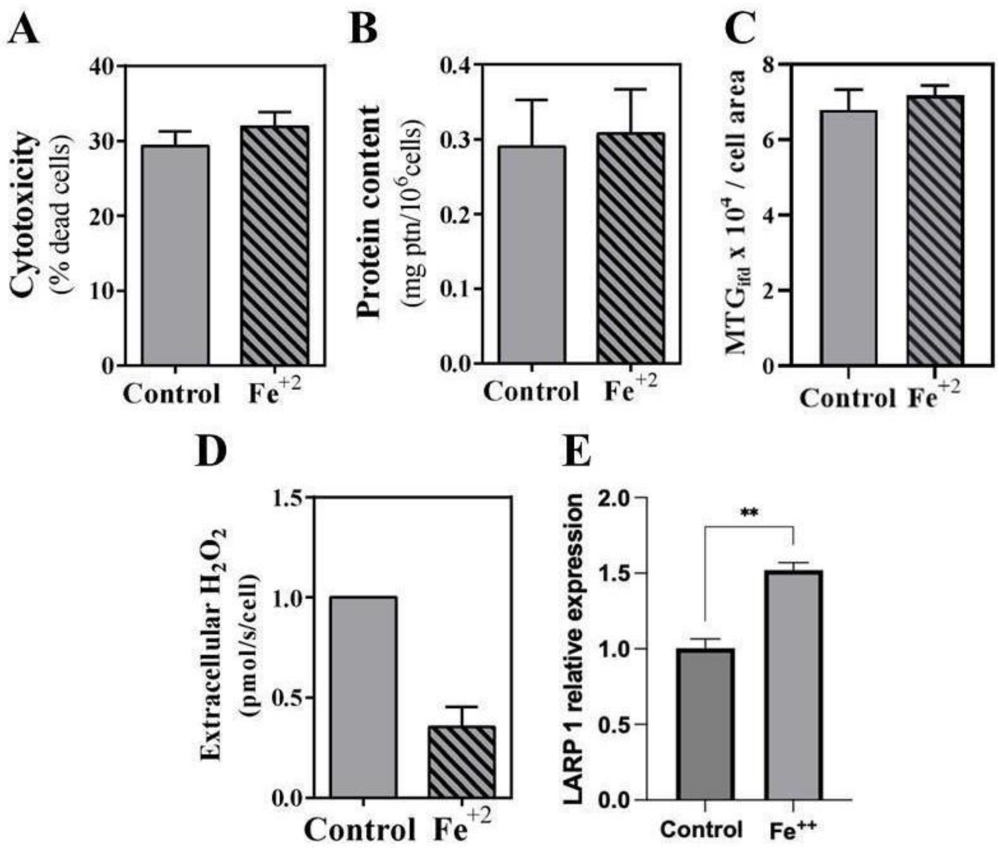


**Fig. S1** Additional characterization of neuroblastoma cells exposed to 100 µM iron for 24 h. (A) Cytotoxicity assessed by extracellular LDH release. (B) Total protein content determined by the Lowry method. (C) Mitochondrial content expressed as MTG integrated fluorescence density per cell area. (D) Extracellular $H_2O_2$ production, quantified by Amplex Red oxidation and fluorescence spectroscopy. (E) *LARP1* mRNA relative expression determined by RT-qPCR ($2^{-\Delta\Delta Ct}$ method, *GUSB* as reference gene). Grey bars denote control and hatched bars denote 100 µM iron-treated cells. Data are expressed as the mean ± SEM of four independent experiments. $**p<0.01$ versus control
